# Investigating the antimicrobial and immunomodulatory effects of honeybee venom peptide apamin in the *Drosophila* genetic platform

**DOI:** 10.1101/2024.07.25.605111

**Authors:** Yanan Wei, Wenjie Jia, Yanying Sun, Tianmu Zhang, Hongyu Miao, Zekun Wu, Ran Dong, Fangyong Ning, Woo Jae Kim

## Abstract

Apamin, an 18-amino acid peptide neurotoxin, constitutes a significant portion of honeybee venom. Although traditionally recognized for its neurotoxic effects, our research demonstrates that apamin exhibits potent antimicrobial properties when genetically expressed in *Drosophila*. The antimicrobial efficacy of apamin is independent of its disulfide bonds and is enhanced when the peptide is membrane-tethered. This expression selectively targets and inhibits specific harmful bacterial species, such as *Pseudomonas aeruginosa*, *Enterococcus faecalis*, and *Escherichia coli*, while promoting beneficial bacteria like *Lactobacillus plantarum* thereby improving the gut microbiome. The antimicrobial activity of apamin is localized to the gut and is associated with increased proliferation of intestinal stem cells, acidification of the midgut pH, and activation of enteroendocrine cell calcium signaling. Furthermore, apamin’s antimicrobial function is dependent on specific peptidoglycan recognition proteins, with PGRP-LA and PGRP-SCs being essential. Apamin expression alone is sufficient to restore the integrity of the gut barrier compromised by stressed conditions. Ultimately, apamin supplementation enhances honeybee gut health, particularly in the presence of ingested bacteria. The expression of other honeybee antimicrobial peptides also significantly reduces bacterial infection in flies. Overall, our study provides a comprehensive understanding of the molecular function and regulation of honeybee venom peptides and antimicrobial peptides, utilizing the *Drosophila* model system to unravel their mechanisms of action and therapeutic potential.

## INTRODUCTION

Honeybees, specifically *Apis mellifera* and *Apis cerana*, are vital for pollinating many crops and have a significant impact on the general stability of ecosystems ^1^.

Unfortunately, honeybee populations are currently facing a decline, primarily due to a range of threats including pests, pathogens, genetic bottlenecks, and environmental challenges. This decline in honeybee populations poses a significant risk to both agriculture and the environment ^2^.

Antimicrobial peptides (AMPs) and other peptides produced by honeybees play crucial roles in their innate immune defense mechanisms. These peptides can effectively target and eliminate a wide range of pathogens, making them promising candidates for the development of novel genetic strategies to improve honeybee immunity ^3^. However, the molecular mechanisms underlying the function and regulation of these peptides remain largely unknown.

Venom peptides (VPs), synthesized by a broad range of organisms including honeybees, have attracted significant interest due to their diverse and potent biological effects. VPs have been found to exhibit a range of functions, including antimicrobial, antiviral, and anticancer properties, making them promising candidates for the development of novel therapeutic agents ^4^. In particular, honeybee VPs have been a subject of intense research, as they possess unique properties that make them highly valuable for medical purposes ^5,6^.

Apamin, an 18 amino acid peptide neurotoxin, is one of the bioactive components of bee venom, making up 2%–3% of its total dry weight ^7–10^. Apamin is a specific inhibitor of small conductance calcium-activated potassium (SK) channels, which are critical in various diseases ^11–19^. Despite these advances, the molecular mechanisms and pathogenesis of SK channel blockers and their anti-inflammatory effects are not fully understood.

The evolutionary proximity between apamin-producing honeybees and the fruit fly, *Drosophila melanogaster*, suggests a conserved biochemical and genetic foundation ^20^. This conservation presents an avenue to exploit the sophisticated genetic tools available in *Drosophila* to unravel the intricate molecular mechanisms underlying the function of apamin ^21,22^. Utilizing powerful genetic resources available in fruit fly, we are able to delineate the precise molecular pathways by which apamin mediates its biological effects ^23–25^. The genetic analysis of apamin within the *Drosophila* model system holds substantial promise for advancing our understanding of this peptide’s mechanisms of action and for enhancing its therapeutic potential.

## RESULTS

### Genetically encoded apamin has antimicrobial peptide activity regardless of its disulfide bridge formation

To assess the functionality of genetically encoded honeybee VPs in the *Drosophila* model, we developed *UAS-Melittin* and *UAS-Apamin* constructs that incorporate a previously characterized signal peptide at their N-termini ^26^ (Fig. 1a). Despite the extensive literature on melittin’s antimicrobial properties ^27^, its broad expression in flies did not diminish the bacterial infection by *Pseudomonas aeruginosa*, a gram-negative pathogen that commonly afflicts humans, insects, and plants ^28,29^ (Fig. 1b and Fig. S1b-d). In contrast, apamin, a neurotoxin known for its neuronal effects ^11^, displayed unexpected antimicrobial activity when expressed genetically in flies (Fig. 1c). These findings suggest that while melittin did not manifest its antimicrobial function when genetically introduced, apamin demonstrated potential as an AMPs when encoded genetically.

**Fig 1:**
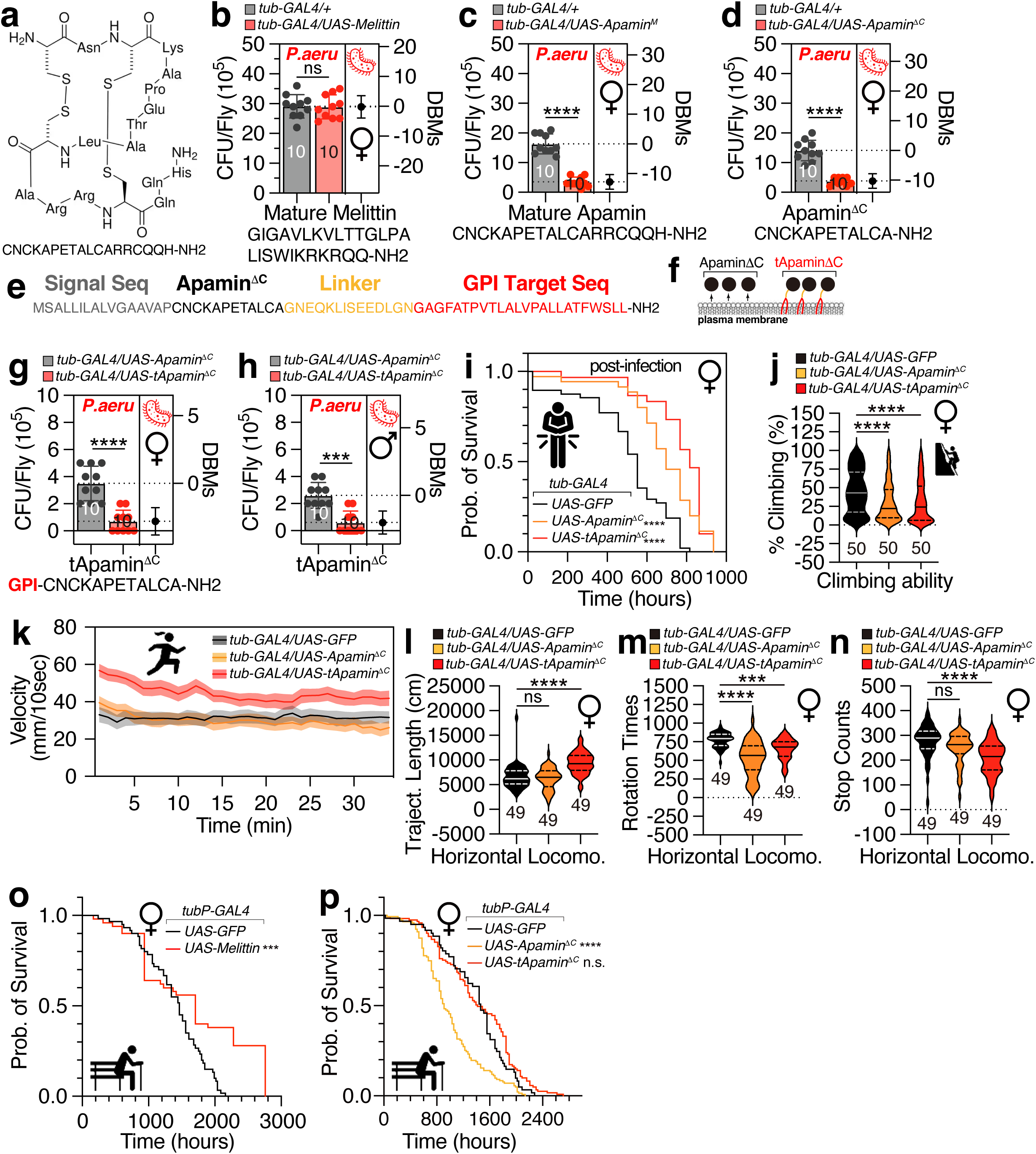
The structure and sequences of apamin related amino acids, the antimicrobial effect, survival curve and locomotion of flies expressing different honeybee VPs. **a,** Chemical structure of mature apamin. **b,** Infectious dose of female flies expressing mature format melittin by *tub-GAL4* following a 12-hour orally feeding of *P. aeruginosa* culture. CFU stands for Colony-Forming Unit. DBMs represent difference between means, which is a statistical measure that quantifies the average discrepancy between two groups. Numbers shown are sample sizes for each condition. (n=10) **c,** Infectious dose of female flies expressing mature format apamin by *tub-GAL4*. **d,** Infectious dose of female flies expressing Apamin^ΔC^ by *tub-GAL4*. (n=10) **e,** The amino acid sequence of tethered form Apamin^ΔC^, including signal sequence (grey), Apamin^ΔC^ sequence (black), linker sequence (yellow), GPI target sequence (orange). **f,** The diagram of Apamin^ΔC^ and tethered form of Apamin^ΔC^ (tApamin^ΔC^). **g,** Infectious dose of female flies and **h,** male flies expressing Apamin^ΔC^ and tApamin^ΔC^ by *tub-GAL4*. **i,** The survival curve of female Apamin^ΔC^ and tApamin^ΔC^ flies post infection by *P. aeruginosa* culture. **j,** The climbing ability, **k,** velocity, **l,** trajectory length, **m,** rotation times, and **n,** stop counts of female Apamin^ΔC^ and tApamin^ΔC^ flies (n=49 for each condition). **o,** The survival curve of female flies expressing melittin via *tub-GAL4* (n=50 for each condition). **p,** The survival curve of female flies expressing Apamin^ΔC^ and tApamin^ΔC^ via by *tub-GAL4* driver (n=50 for each condition).

To assess whether the antimicrobial function of apamin is dependent on its disulfide bridges (Fig. S1a), we engineered *UAS-Apamin*^Δ*C*^, a variant lacking six carboxyl-terminal residues and found that Apamin^ΔC^ retained antimicrobial activity comparable to the full-length peptide (Fig. 1d and Fig. S1e). The efficacy of genetically expressed apamin was consistent across various GAL4 drivers, indicating that the strength of GAL4 is not a critical factor for apamin’s action (Fig. S1f).

The membrane tethering of AMPs can enhance their activity, but this modification is challenging to achieve under *in vitro* conditions ^30^. We addressed this challenge by genetically encoding apamin to be covalently linked to a glycosylphosphatidylinositol (GPI) anchor on the extracellular leaflet of the plasma membrane (tApamin^ΔC^) ^26^ (Fig. 1e-f). This membrane-tethered apamin exhibited a significant increase in antimicrobial effect (Fig. 1g-h) and prolonged survival following infection in female flies (Fig. 1i). Although the broad expression of secreted or membrane-tethered apamin slightly reduced climbing ability (Fig. 1j), it increased locomotor behaviors such as forward velocity (Fig. 1k), trajectory length (Fig. 1l), and trajectory percentage (Fig. S1g) while decreasing rotation times (Fig. 1m) and stop counts (Fig. 1n). The lifespan of flies expressing melittin was significantly reduced (Fig. 1o), in contrast to apamin expression, which did not affect the lifespan of female flies or have slight effect on male flies (Fig. 1p and Fig. S1h). The collective data suggest that the widespread expression of membrane-tethered apamin in flies confers potent antimicrobial activity, alters locomotor behaviors, and does not impact the lifespan of the flies.

### Apamin expression alters the composition of the gut microbiome environment

To evaluate the efficacy of genetically expressed apamin against various bacterial infections, we infected fruit flies with Gram-positive *Enterococcus faecalis*, a common intestinal pathogen, and observed that apamin expression effectively inhibited *E. faecalis* infection (Fig. 2a). Additionally, apamin expression was found to completely block *E. coli* infection (Fig. 2b). However, in contrast to *E. faecalis* and *E. coli*, apamin expression did not reduce the infection caused by *Lactobacillus plantarum*, a bacterium commonly found in fermented foods and used as a probiotic (Fig. 2c). Furthermore, apamin expression did not reduce the infection caused by *Apibacter raozihei*, which are present in the guts of *Apis* species ^31^ (Fig. 2d).

**Fig 2:**
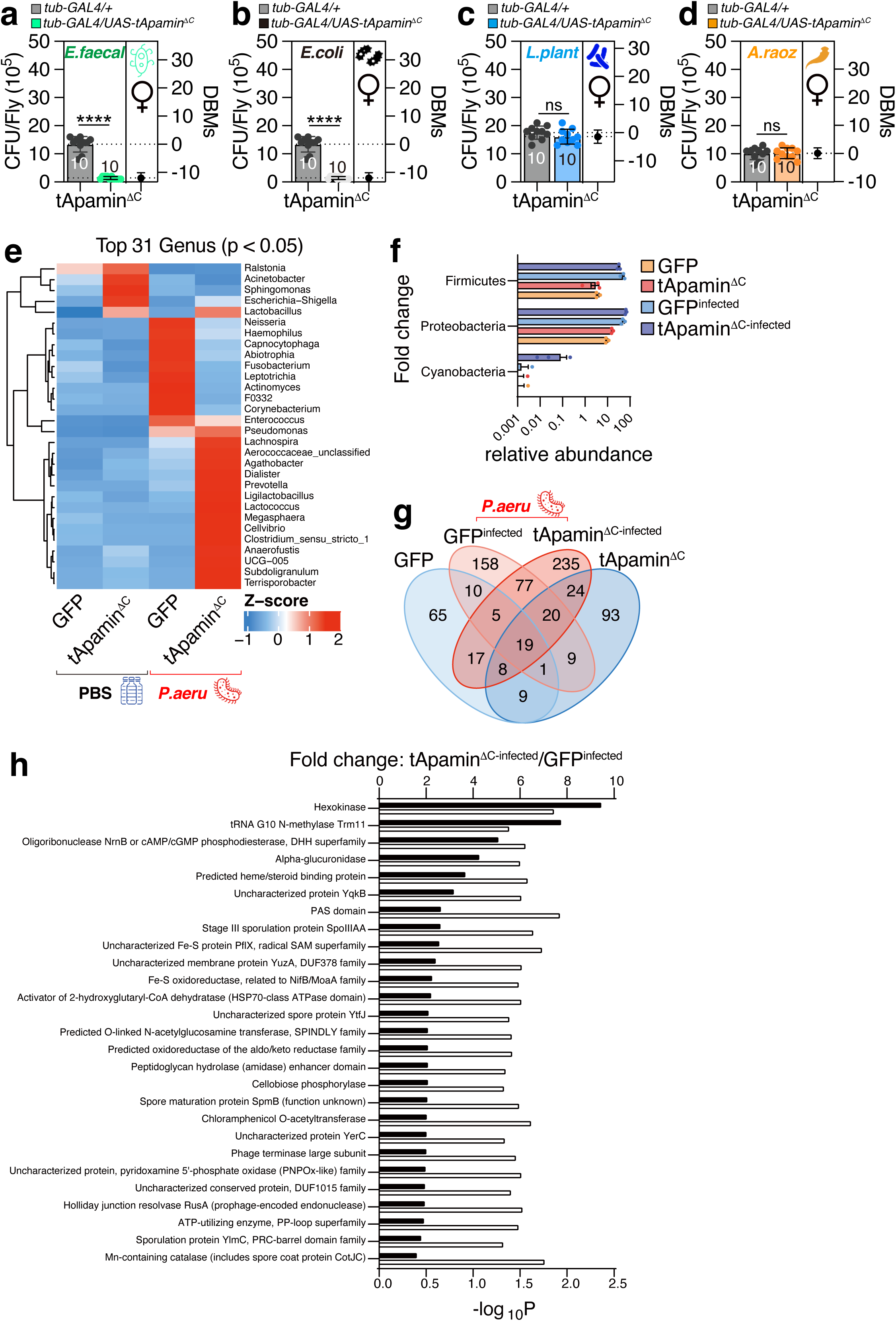
Infectious dose of flies expressing tApamin^ΔC^ encountering different bacteria and 16S rRNA sequencing results. **a,** Infectious dose of female flies expressing tApamin^ΔC^ by *tub-GAL4* following a 12-hour orally feeding of *E. faecalis* culture, **b,** *E. coli* culture, **c,** *Lactobacillus plantarum* culture, and **d,** *Apibacter raozihei* culture. **e,** 16S rRNA sequencing abundance heatmaps of top candidate genus (P value < 0.05, in the comparison of infected and uninfected, tApamin^ΔC^ versus control; n = 3 for each condition). Heatmaps show Z-scores of relative abundances. Rows were ordered by hierarchical clustering. Data from ≥ 3 biological replicates per condition. **f,** Relative abundance of the top candidate Phyla (P value < 0.05, in the comparison of infected and uninfected, tApamin^ΔC^ versus control; n = 3 for each condition). **g,** Venn diagram based on the Amplicon Sequence Variants (ASVs) observed in different groups. **h,** Bacterial COG functional category prediction by PICRUSt2. The figure demonstrates the COG functional categories. Black bar: Foldchange of relative abundance; white bar: −log_10_ P values for each COG term.

To validate the efficacy of our infection protocols and to evaluate the impact of apamin expression on gut microbiota, we conducted 16S ribosomal RNA (rRNA) sequencing (Fig. S2a). We confirmed that *Pseudomonas aeruginosa* infection markedly increased the abundance of *Pseudomonas* genus within the fly gut (Fig. S2b). In uninfected flies, apamin expression increased the relative abundance of *Ralstonia*, *Acinetobacter*, *Sphingomonas*, *Escherichia-Shigella*, and *Lactobacillus* (left two panels of Fig. 2e). Remarkably, membrane-tethered apamin expression fundamentally transformed the gut microbiome composition in flies infected with *P. aeruginosa* (right two panels of Fig. 2e). Specifically, the abundance of *Lachnospira*, known for its role in early infant gut health ^32^, and *Prevotella*, a microbe crucial for human health and disease balance ^33^, increased. Additionally, *Lactobacillus, Lactococcus, Subdoligranulum*, commonly used as probiotics in humans ^34–36^, and *Megasphaera*, which has unique roles in reproductive health ^37^ were enriched (Fig. 2e). The most notable increase was in the *Cyanobacteria* phylum, a group of autotrophic, gram-negative bacteria capable of oxygenic photosynthesis ^38^ (Fig. 2f). The appearance of approximately 235 new ASVs (amplicon sequence variants) upon apamin expression suggests that apamin influences both the commonalities and divergences within the gut bacterial community (Fig. 2g).

Bacterial COG (Clusters of Orthologous Genes) functional category analysis revealed an increase in hexokinase function upon *P. aeruginosa* infection with apamin expression. Hexokinase has been identified as an innate immune receptor for bacterial peptidoglycan detection ^39^, indicating that apamin expression enhances the bacterial community for innate immunity (Fig. 2h). Bacterial KEGG (Kyoto Encyclopedia of Genes and Genomes) pathway analysis indicated that xylene degradation and primary bile acid biosynthesis were slightly reduced after infection with apamin expression (Fig. S2c). Collectively, these findings indicate that the genetic expression of apamin selectively targets and inhibits specific bacterial species, predominantly harmful bacteria, while promoting the beneficial ones, thereby enhancing the gut microbiome community.

### Genetic expression of apamin beneficially changes gut environment

To investigate the specific tissue in which apamin exerts its antimicrobial effects, we utilized various GAL4 drivers targeting specific tissues to express apamin. Our results indicated that the expression of apamin in either enteroendocrine cells (EEs) or intestinal stem cells (ISCs) was sufficient to recapitulate the effects observed with tub-GAL4 expression (Fig. 3a-i and Fig. S3a-j), suggesting that EEs or ISCs are pivotal in apamin’s ability to eliminate harmful bacteria within the fly gut.

**Fig 3:**
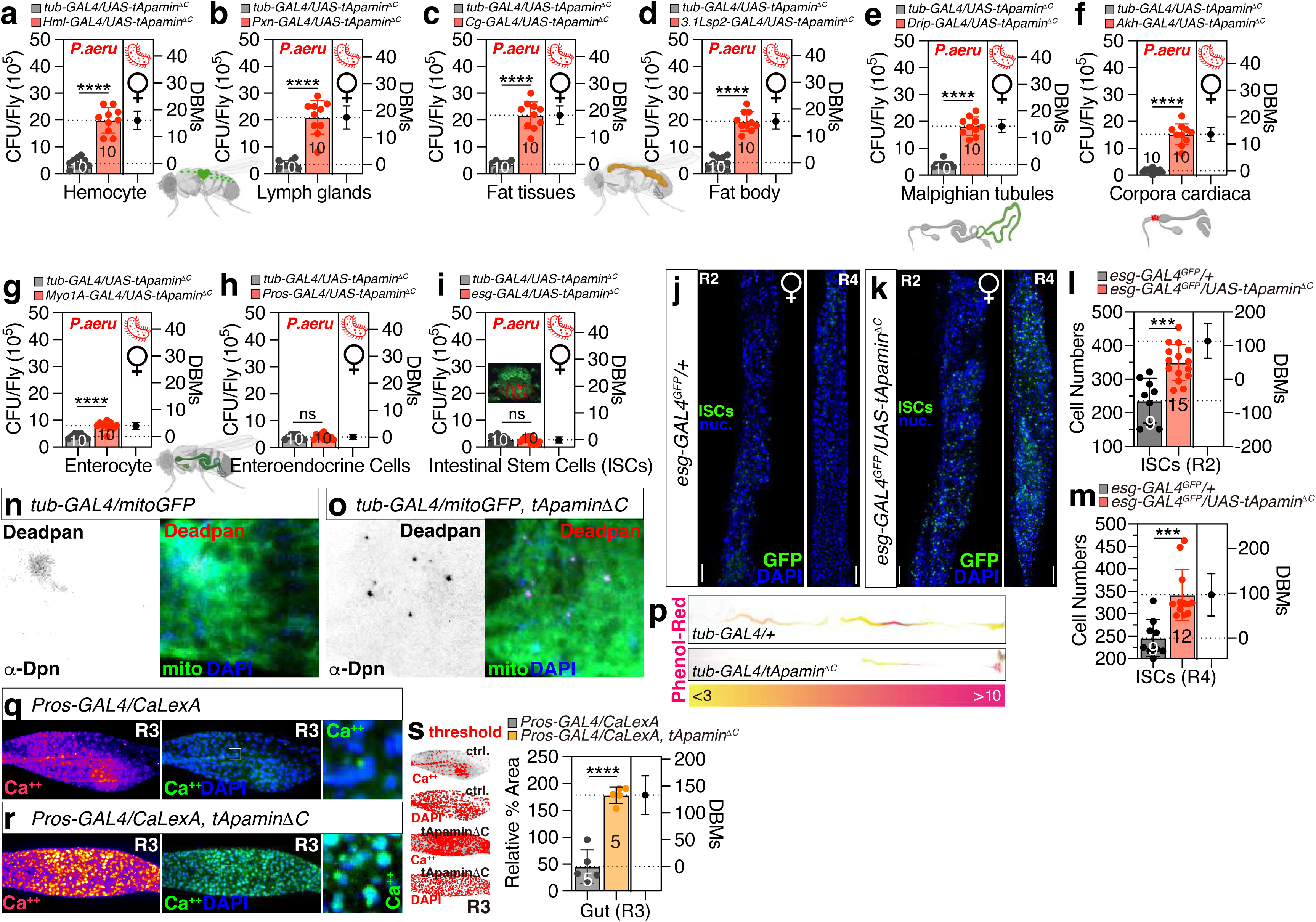
Tissue screening of antimicrobial effects in tApamin^ΔC^ expressing female flies and gut environment changes displayed. **a-i,** Infectious dose between female flies expressing tApamin^ΔC^ via a, *tub-GAL4* and *Hml-Gal4*, b, *Pxn-GAL4*, c, *Cg-GAL4*, d, *3.1Lsp2-GAL4*, e, *Drip-GAL4*, f, *Akh-GAL4*, g, *Myo1A-GAL4*, h, *Pros-GAL4*, i, *esg-GAL4*, following a 12-hour orally feeding of *P. aeruginosa* culture. **j, k,** The ISCs in R2 and R4 regions between normal condition and tApamin^ΔC^ expression female flies driven by *esg-GAL4* with *UAS-GFP*, fly guts were immunostained with anti-GFP (green), and DAPI (blue). Scalebar 200µm. **l, m,** The ISC number counts in R2 region **(l)** and R4 region **(m)** between normal condition (n=9) **(j)** and tApamin^ΔC^ expressing female flies (n=15 for R2 region, n=12 for R4 region) **(k)**. **n,o,** The Dpn-positive neuroblast cells and mitochondria (*UAS-mitoGFP*) in midgut of female flies in normal (n=3) **(n)** and tApamin^ΔC^ expressing conditions (n=3) **(o)** by *tub-GAL4*, fly guts were immunostained with anti-Deadpan (red), anti-GFP (green), and DAPI (blue). Scalebar 200µm. **p,** The pH zones in the full gut of female flies in normal and tApamin^ΔC^ expressing conditions via *tub-GAL4* driver (n=5), flies were fed by phenol red dye which changes from yellow at pH<3, an acidic region, to brighter red at pH>10, an alkaline region. **q-s,** CalexA assay for *Pros-GAL4* together with *lexAop-mCD8GFP; UAS-CaLexA, lexAop-CD2-GFP* of control (n=5) **(q)** and tApamin^ΔC^ expressed (n=5) **(r)** female flies at R3 region of gut, and were immunostained with anti-GFP (green), and DAPI (blue). The thresholds were adjusted and calcium signal was quantified as relative percentage area normalized by DAPI signal **(s)**.

Furthermore, we observed that the expression of membrane-tethered apamin in ISCs led to a significant increase in stem cell numbers in both the R2 and R4 regions of the fly gut (Fig. 3j-m and Fig. S3k-l), indicating that apamin may promote stem cell proliferation. Additionally, apamin expression increased the number of Dpn-positive neuroblast cells in the midgut (Fig. 3n-o) without altering mitochondrial numbers or activity (Fig. S3m-p). Since Dpn enhances the self-renewal capacity of stem cell populations ^40,41^, these data suggest that apamin can promote the self-renewal of ISCs through a noncanonical pathway.

Moreover, the expression of membrane-tethered apamin in EEs led to a significant change in the pH of the midgut, making it more acidic, as evidenced by phenol-red dye absorption (Fig. 3p). However, apamin expression did not alter the level of reactive oxygen species (ROS) in the midgut (Fig. S3q-r). Surprisingly, the genetic expression of membrane-tethered apamin in EEs resulted in a dramatic increase in calcium levels in the R3 region of the gut (Fig. 3q-s). Therefore, the potent antimicrobial activity of membrane-tethered apamin appears to be attributed to the enhanced proliferation of ISCs, the acidification of the midgut pH, and the activation of EE calcium signaling.

### Apamin expression exerted nuanced effects on neuronal behaviors yet demonstrated potential to mitigate pathological responses induced by stress

Apamin is known as a potential neurotoxin that acts as a blocker of small conductance calcium-activated potassium (SK) channels ^11^. We investigated its impact on nervous system function. Neuronal expression of apamin slightly reduced forward velocity (Fig. S4a), while trajectory length and area were comparable to controls (Fig. S4a-b). Flies expressing apamin exhibited fewer turns, but similar stop frequencies compared to controls (Fig. S4c-e).

Interestingly, flies expressing membrane-tethered apamin in neurons exhibited reduced movement speed (Fig. 4a), distance (Fig. 4b-c), and turns (Fig. 4d), but normal stopping behavior (Fig. 4e). This suggests that neuronal expression of membrane-tethered apamin has the opposite effect on locomotion compared to whole-body expression (Fig. 1k vs. Fig. 4a).

**Fig 4:**
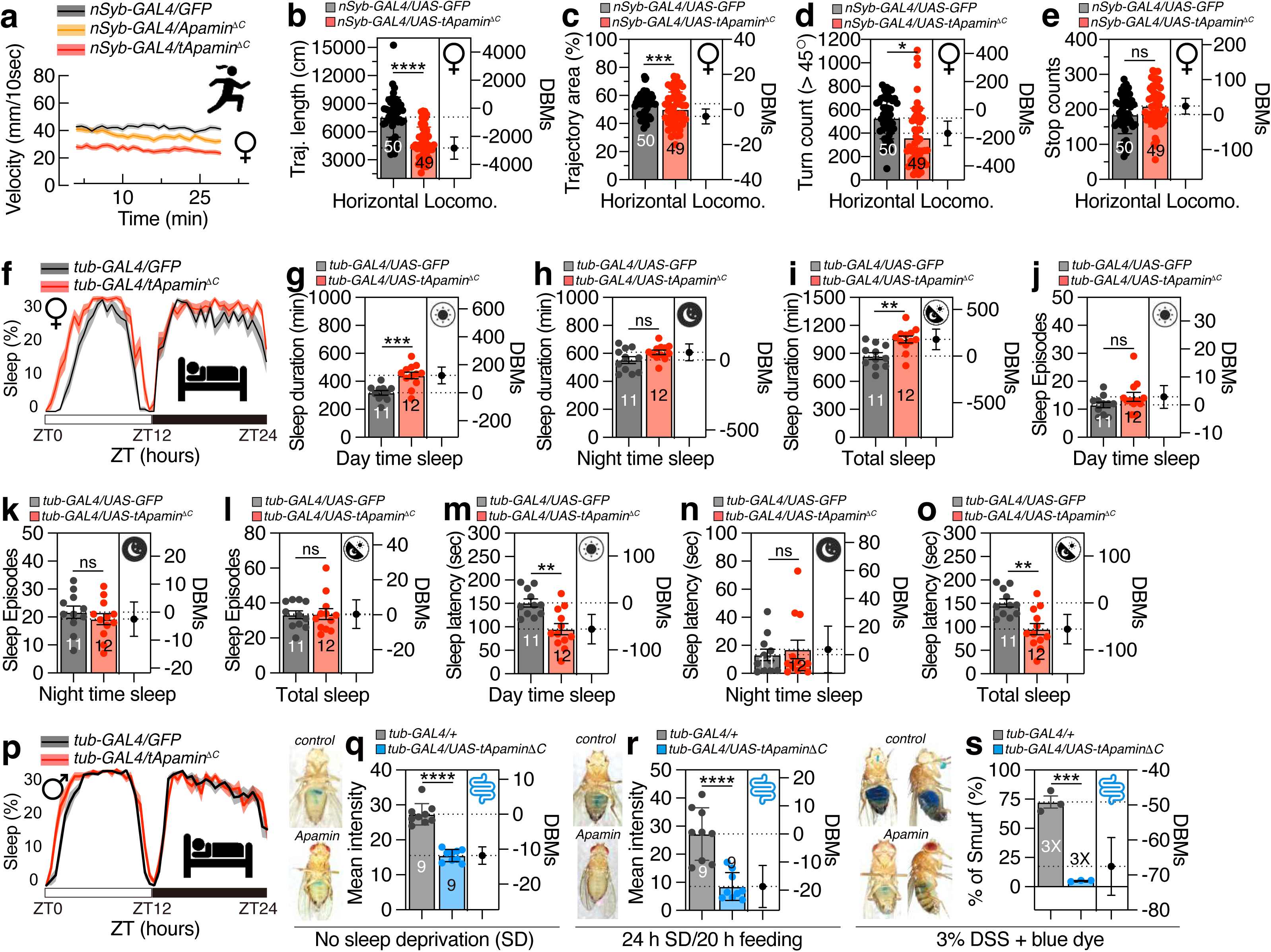
Locomotion of neuronal tApamin^ΔC^ expression fly, sleep pattern of pan-expressed tApamin^ΔC^ fly and Smurf results induced by different stress conditions. **a,** The velocity, **b,** trajectory length, **c,** trajectory area, **d,** rotation times, and **e,** stop counts of female flies expressing tApamin^ΔC^ by *nSyb-GAL4* (n=49)compared with controls (n=50). **f-p,** Sleep profiles (average proportion of time spent sleeping in consecutive 30-min segments during a 24-h LD cycle) and quantification of female flies **(f)** and male flies **(p)** expressing tApamin^ΔC^ expression by *tub-GAL4* driver. Quantification of sleep episodes for tApamin^ΔC^ expression flies (n=11 for controls, n=12 for tApamin^ΔC^ expressing flies) **(g-i)**. Quantification of sleep episodes for tApamin^ΔC^ expression flies **(j-l)**. Quantification of sleep latency for tApamin^ΔC^ expressing flies **(m-o)**. **q,** The mean intensity of Smurf assay flies expressing tApamin^ΔC^ compared with controls under normal condition. The experiment was repeated three times and one representative figure was shown of each condition. **r,** The mean intensity of Smurf assay flies expressing tApamin^ΔC^ compared with controls (n=9) after 24 hours sleep deprivation and 20 hours dye feeding. The experiment was repeated three times and one representative figure was shown of each condition. **t,** The percentage of flies (n=20, repeated for 3 times) showing Smurf phenotype after a 3% DSS feeding with blue dye. The experiment was repeated three times and one representative figure was shown of each condition.

It is well-established that potassium channels play a crucial role in sleep modulation in both mammals and flies ^42–49^. In mice, apamin treatment has been shown to suppress rapid eye movement (REM) sleep without causing a compensatory rebound^50^. Expression of the secreted form of apamin increased sleep duration and daytime sleep episodes while reducing sleep latency (Fig. S4f-o), indicating a distinct effect on sleep in fly neurons compared to mammalian nervous systems. Expression of the membrane-tethered form of apamin slightly increased daytime sleep duration and decreased daytime sleep latency without affecting sleep episodes in males and females (Fig. 4f-p), suggesting a contrasting effect of apamin on sleep compared to vertebrates.

Sleep deprivation can lead to mortality through the accumulation of reactive oxygen species (ROS) in the gut, and gut neuropeptides mediate energy depletion induced by sleep loss in *Drosophila* ^51,52^. Sleep deprivation elicits perturbations in the intestinal epithelial barrier integrity, a phenomenon that represents an evolutionarily preserved feature of senescence and is associated with alterations in metabolic and inflammatory biomarkers ^53,54^. Our findings indicate that apamin expression can mitigate gut barrier dysfunction induced by sleep deprivation ^55^ (Fig. 4q-s and Fig. S4p). Consequently, the presence of apamin in the gastrointestinal tract of *Drosophila* can serve as a protective mechanism against ROS-mediated stress induced by sleep loss.

Administration of dextran sulfate sodium (DSS) induces mucosal injury in the adult *Drosophila* gastrointestinal tract, subsequently impacting viability ^56,57^. The DSS-mediated intestinal inflammation model has gained widespread recognition as an effective experimental paradigm for colitis and inflammatory bowel disease (IBD) research ^58,59^. Our findings demonstrate that the expression of apamin is capable of ameliorating DSS-triggered intestinal inflammation (Fig. 4t and Fig. S4q-r), suggesting that apamin functions as a potent inhibitor of DSS-induced epithelial damage.

### Specific peptidoglycan recognition proteins (PGRPs) are required for antimicrobial function of apamin

Peptidoglycan recognition proteins (PGRPs) are pivotal in the detection and response to peptidoglycan, a fundamental component of bacterial cell walls. The recognition of peptidoglycan by PGRPs initiates a signaling cascade that activates immune responses, including the Toll and IMD signaling pathways, which, in turn, induce the production of AMPs by the transcription factor Relish (Rel) ^60,61^. In *Drosophila*, the PGRP family comprises various members, including PGRP-L (long forms), PGRP-S (short forms), and proteins that are transmembrane, intracytoplasmic, or secreted.

These proteins are present in various tissues and cell types, such as hemocytes, epithelial cells, and fat body cells, and are indispensable for the immune system’s proper functioning ^62–64^ (Fig. 5a).

**Fig 5:**
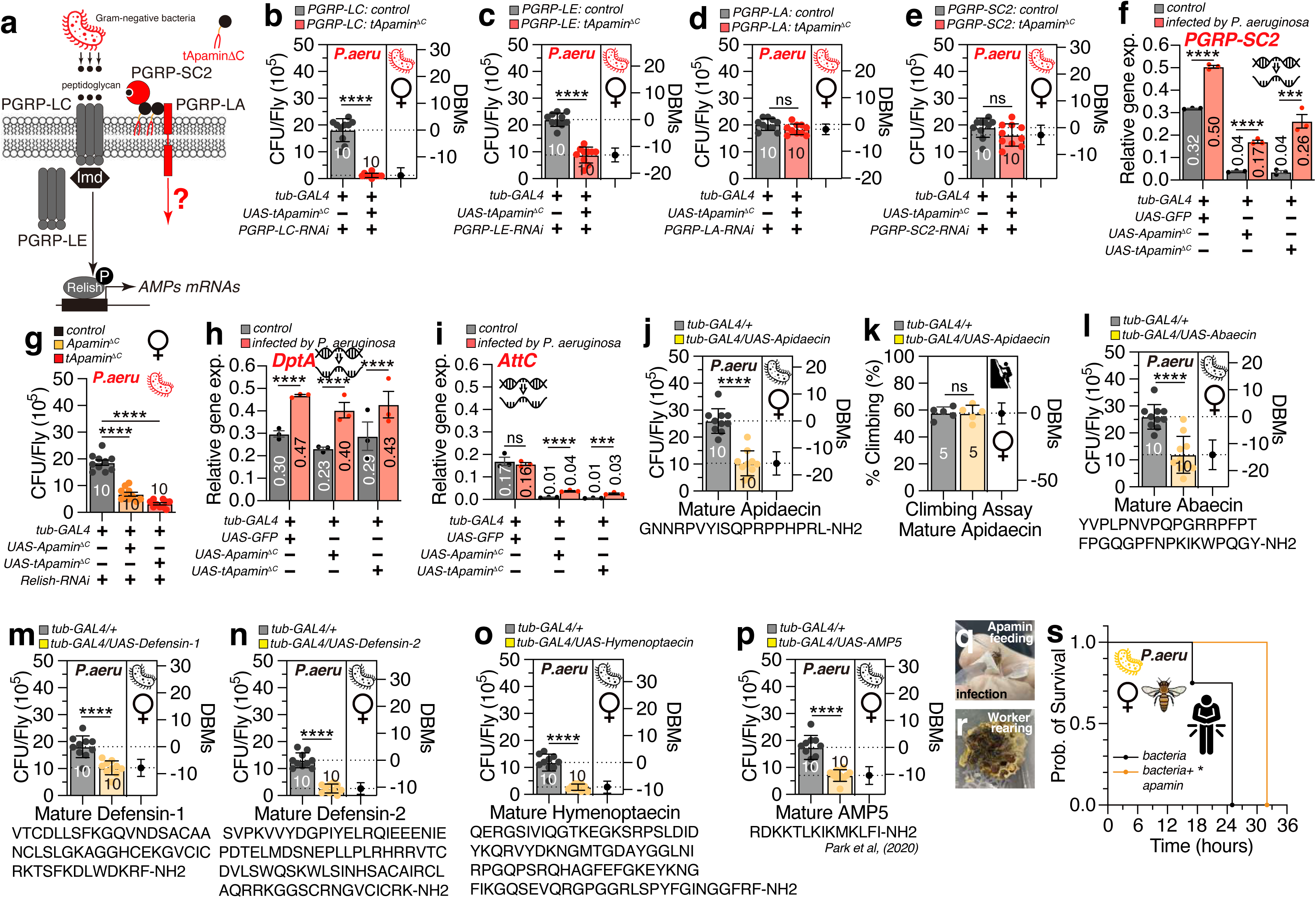
Possible immune pathways and AMPs related and apamin application to honeybees. **a,** The diagram of tApamin^ΔC^ involved in PGRP family related immune pathway. **b-e,** Infectious dose of controls and female flies expressing tApamin^ΔC^ via *tub-GAL4*, with a knockdown of **b,** *PGRP-LC*, **c,** *PGRP-LE*, **d,** *PGRP-LA*, **e,** *PGRP-SC2*, following a 12-hour orally feeding of *P. aeruginosa* culture. (n=10) **f,** *PGRG-SC2* relative expression level between control, Apamin^ΔC^ and tApamin^ΔC^ expression flies, under *P. aeruginosa* infected or uninfected conditions respectively (repeated for 3 times). **g,** Infectious dose of female flies expressing Apamin^ΔC^ and tApamin^ΔC^ by *tub-GAL4*, with a knockdown of *Relish* compared with control. (n=10) **h,** *DptA* and **i,** *AttC* relative expression level between control, Apamin^ΔC^ and tApamin^ΔC^ expression flies, under infected or uninfected conditions respectively (repeated for 3 times). **j,** Infectious dose and **k,** Climbing ability of controls and female flies expressing Apidaecin by *tub-GAL4*. (n=5) **l-p,** Infectious dose of controls and female flies expressing Abaecin **(l)**, Defendin-1 **(m)**, Defensin-2 **(n)**, Hymenoptaecin **(o)**, and AMP5 **(p)** by *tub-GAL4* following a 12-hour orally feeding of *P. aeruginosa* culture. (n=10) **q,** Honeybee apamin feeding and infection assay application. **r,** Worker bee rearing. **s,** The survival curve of honeybees infected orally by *P. aeruginosa*, control (black) (n=4) compared with high concentration apamin (yellow) administration (n=2).

Knockdown of PGRP-LC or LE did not impact the antimicrobial effect of apamin (Fig. 5a-b), indicating that the antimicrobial action of apamin does not rely on the function of PGRP-LC and LE (Fig. 5a). Conversely, knockdown of PGRP-LA or SC2 abolished the antimicrobial effect of apamin (Fig. 5d-e), suggesting that apamin’s antimicrobial activity depends on the function of PGRP-LA and SC2 to resist bacterial infection. Expression of apamin significantly reduced PGRP-SC2 levels in the uninfected fly gut, while bacterial infection increased PGRP-SC2 expression as observed in control flies (Fig. 5f), indicating that apamin can normalize PGRP-SC2 expression by improving gut microbiota under non-infected conditions but does not inhibit infection-induced PGRP-SC2 expression.

Relish, a transcription factor, is a crucial downstream component of the IMD pathway, which governs the antibacterial response ^61^. Knockdown of Rel did not diminish the antimicrobial activity of either the secreted or membrane-tethered form of apamin (Fig. 5g and Fig. S5a), suggesting that Rel-mediated transcriptional regulation is not necessary for apamin’s antimicrobial properties. Furthermore, apamin expression did not affect the expression of AMPs like DptA and AttC (Fig. 5h-i), indicating that apamin’s antibacterial function is independent of Rel-mediated transcriptional activation of AMP genes.

Honeybees allocate significant energetic and physiological resources to immunity to counterbalance the risks of disease associated with their social lifestyle, and AMPs are integral to their innate immune defenses ^3,65^. Seven AMPs have been identified in honeybee hemolymph, encoded by a total of 19 cDNAs ^3^, but the molecular mechanisms governing honeybee AMPs are not well understood due to the lack of genetic tools. To gain a more detailed understanding of the molecular function of honeybee AMPs, we developed fly strains that express all honeybee AMPs using the UAS cassette. We discovered that the expression of honeybee AMPs in the fruit fly platform can significantly reduce bacterial infection without affecting climbing ability in both males and females (Fig. 5j-o and Fig. S5b-i). Additionally, the recently identified uncharacterized honeybee AMP5 also demonstrated robust antibacterial activity in the fly system (Fig. 5p and Fig. S5j) ^66^, suggesting that the molecular mechanisms of honeybee AMPs can be elucidated in detail using the evolutionary proximate *Drosophila* platform.

### The administration of apamin to honeybees enhance survival rates following bacterial infection

To assess the applicability of apamin’s protective effects observed in the *Drosophila* model to honeybee physiology, a series of experiments were conducted involving the feeding of apamin to honeybees (Fig. 5q-r and Movie. S1-2). The dietary introduction of apamin exerted no adverse effects on honeybee viability. Indeed, the administration of a low dose of apamin significantly increased the lifespan of worker honeybees isolated from their colony (Fig. S5k), indicating that apamin supplementation may confer survival benefits to honeybees. Moreover, apamin supplementation was found to increase the survival rate of honeybee workers challenged with a substantial bacterial load (Fig. 5s). These findings indicate that apamin exhibits a protective effect against bacterial infections in the honeybee gut, akin to the results observed in the *Drosophila* model.

## DISCUSSION

Utilizing the fruit fly model system, we successfully demonstrated the expression of honeybee venom peptide, apamin in *Drosophila* displaying unexpected antimicrobial activity. The antimicrobial function of apamin is shown to be independent of its disulfide bridges and is enhanced when the peptide is membrane-tethered (Fig. 1). Notably, apamin expression selectively targets and inhibits specific bacterial species, predominantly harmful bacteria, while promoting beneficial ones, thereby enhancing the gut microbiome community (Fig. 2). Further analysis revealed that apamin’s antimicrobial activity is localized to the gut and is associated with increased proliferation of intestinal stem cells, acidification of the midgut pH, and activation of enteroendocrine cell calcium signaling (Fig. 3). Additionally, we explored the effects of apamin on nervous system function, finding that neuronal expression of membrane-tethered apamin mildly impairs locomotion and sleep (Fig. 4). Importantly, the antimicrobial function of apamin is found to depend on specific peptidoglycan recognition proteins (PGRPs), with PGRP-LA and SC2 being essential. Furthermore, we extended our findings to other honeybee AMPs, demonstrating their potent antimicrobial activity in the fly system (Fig. 5). Overall, our research provides a comprehensive understanding of the molecular function and regulation of honeybee VPs and AMPs, utilizing the *Drosophila* model system to unravel their intricate mechanisms of action and therapeutic potential.

The expression of apamin significantly modulates the physiological landscape of the fruit fly gut (Fig. 3 and Fig. S3). A particularly intriguing observation is that apamin has the capacity to stimulate the proliferation of intestinal stem cells (ISCs). Our findings indicate a marked increase in the number of Dpn-positive neuroblast-like cells in the presence of apamin, which is consistent with the role of Dpn as a neural stem cell factor that promotes self-renewal and inhibits differentiation into EE cells ^67^. This leads us to hypothesize that apamin may phenocopy the effects of Ttk depletion, which can initiate cell transdifferentiation from differentiated ECs to EE-like cells by derepressing neuroblast-specific transcription factors such as Dpn ^68^. Notably, while Ttk depletion leads to neuroendocrine tumors, apamin expression does not appear to significantly reduce lifespan or locomotor ability, suggesting that it may moderately activate ISC self-renewal without triggering a tumor phenotype.

The physiological roles of *Drosophila* SK channels have been postulated to encompass the modulation of nociception, the enhancement of photoreceptor performance through sensitivity control in the initial visual network, and the regulation of courtship memory ^69–71^. Apamin, a well-established blocker of SK channels in mammals, is known to profoundly influence sleep patterns ^50^. However, our current data align with previous findings in suggesting that neither the expression of apamin nor the disruption of *Drosophila* SK channels significantly alter sleep behavior in fruit flies. The differential effects of apamin on sleep across species suggest a potential selectivity for its action on mammalian SK channels, which is congruent with its evolutionary role as a venom peptide targeted against honeybee predators, such as bears and other mammals preying on honey. This selectivity may be a result of the co-evolutionary arms race between honeybees and their mammalian predators, where apamin’s specificity for mammalian SK channels provides a potent defense mechanism against these particular enemies. The absence of a pronounced neuronal phenotype in *Drosophila* following apamin overexpression supports this notion.

However, to date, there is a lack of direct evidence demonstrating the inhibitory effect of apamin on *Drosophila* SK channels. Further research is required to delineate the precise molecular interactions of apamin with *Drosophila* SK channels, which may provide valuable insights into the unique evolutionary functions of this venom peptide. Further exploration of the molecular mechanisms of apamin in both *Drosophila* and honeybees may unravel the potential therapeutic applications of apamin in bolstering honeybee immunity.

## ACKNOWLEDGEMENT

We are very appreciative to the colleagues who supplied us with several fly strains; Dr. Lihua Jin (Northeast Forestry University, Harbin, China). We would also like to thank Wenjing Li, a PhD student, for their help in using the instruments in the lab. Stocks obtained from the Bloomington Drosophila Stock Center (NIH P40OD018537) were used in this study. Transgenic fly stocks and/or plasmids were obtained from the Vienna Drosophila Resource Center (VDRC, www.vdrc.at). The fly stock was obtained from Korea Drosophila Resource Center. This work was supported by Startup funds from HIT Center for Life Science to WJK.

## AUTHOR CONTRIBUTIONS

**Conceptualization:** Woo Jae Kim.

**Data curation:** Yanan Wei, Wenjie Jia, Ran Dong, Hongyu Miao, Zekun Wu, Woo Jae Kim.

**Formal analysis:** Yanan Wei, Wenjie Jia, Hongyu Miao, Zekun Wu, Woo Jae Kim.

**Funding acquisition:** Woo Jae Kim.

**Investigation:** Woo Jae Kim.

**Methodology:** Fangyong Ning, Woo Jae Kim.

**Project administration:** Woo Jae Kim.

**Resources:** Woo Jae Kim.

**Supervision:** Fangyong Ning, Woo Jae Kim.

**Validation:** Yanan Wei, Woo Jae Kim.

**Visualization:** Yanan Wei, Wenjie Jia, Hongyu Miao, Zekun Wu, Woo Jae Kim.

**Writing – original draft:** Woo Jae Kim.

**Writing – review & editing:** Yanan Wei, Yanying Sun, Tianmu Zhang, Hongyu Miao, Woo Jae Kim.

## METHODS

### Fly stocks and husbandry

*Drosophila melanogaster* was raised on cornmeal-yeast medium at similar densities to yield adults with similar body sizes. Flies were kept in 12 h light: 12 h dark cycles (LD) at 25℃ (ZT 0 is the beginning of the light phase, ZT12 beginning of the dark phase). To reduce the variation from genetic background, all flies were backcrossed for at least 3 generations to CS strain. All mutants and transgenic lines used here and their sources were as follows: *w^1118^* (Vienna Drosophila Resource Center, 60000), *w^1118^;UAS-mito-HA-GFP/CyO* (Bloomington Stock Center, 8442), *w^1118^;Cg-GAL4* (Bloomington Stock Center, 7011), *;;tub-GAL4/TM3*, *;;Hml-GAL4*, ;*MyolA-Gal4,UAS-GFP/CyO*, *w*; esg-Gal4,UAS-GFP/CyO* (These four lines were kindly provided by Dr. Lihua Jin, Northeast Forestry University, Harbin, China), *;;3.1Lsp2- GAL4(III)/TM3* (Korea Drosophila Resource Center, 2132), *;; Akh-GAL4* (Bloomington Stock Center, 25684), *y^1^*,*w*;Drip-GAL4/SM6a* (Bloomington Stock Center, 66782), *w*;; pros-GAL4* (Bloomington Stock Center, 80572), *;LexAop- CD8GFP(II);UAS-CaLexA,LexAop-CD2-GFP/TM6B,Tb* (Korea Drosophila Resource Center, 1234), *y^1^,w^1118^;; nSyb-GAL4* (Bloomington Stock Center, 51941), *y^1^*,*sc*,v^1^,sev^21^;; UAS-PGRP-LC-RNAi* (Bloomington Stock Center, 33383), *y^1^*,*sc*,v^1^,sev^21^;UAS-PGRP-LE-RNAi* (Bloomington Stock Center, 60038), *;PGRP- LA-RNAi* (Vienna Drosophila Resource Center, 102277), *y^1^*,*sc*,v^1^,sev^21^;UAS-PGRP- SC2-RNAi* (Bloomington Stock Center, 56915), *y^1^,sc*,v^1^,sev^21^;; Rel-RNAi* (Bloomington Stock Center, 33661).

### Bacteria culture

For *P. aeruginosa* (ATCC 27853), *E. faecalis* (ATCC 29212), *E. coli* (BNCC336902) culture, 10 mL of Luria-Bertani (LB) broth was inoculated with 100 µL of a frozen bacterial stock at 37 °C. And for *L. plantarum* (BNCC336421) and *Apibacter raozihei* (BNCC356061), MRS medium (LABLEAD, 02-293) was used with the same procedure. The main procedure was modified based on previous study^72^. Shake at 150 rpm overnight and grow this subculture in a 1 L conical flask for another night. Pour equal volumes of this subculture across 500 mL centrifuge tubes and spin the subculture at 2,500 x g for 15 min at 4 °C to pellet the bacteria. Remove the supernatant and resuspend the final bacteria pellet in 5% sucrose water solution. Check the OD and adjust to the desired infectious dose (OD600 = 25 in this study).

### Bacterial infection assay

The main procedure was modified based on previous study and described as follows^72^. Flies were starved for 4 h before exposure to bacteria by transferring the flies to empty vials. Place a disc of filter paper on top of food and pipette 100 µL of bacterial culture directly onto the filter disc. For control infections in this study, bacterial culture was replaced with PBS. 5 flies were transferred to the sample tube and leave for 12 h infection exposure. To confirm oral infection, first surface-sterilize the flies immediately after bacterial exposure, by placing them in 100 µL of 70% ethanol for 20–30 s. Remove the ethanol and add 100 µL of triple distilled water for 20–30 s before removing the water. Add 100 µL of 1x PBS and homogenize the fly. Transfer the homogenate to the top row of a 96-well plate and add 90 µL of 1x PBS to every well below. Serially dilute this sample to distinguish a range of CFU values.

Take 10 µL of the homogenate in the top well and add this to the well below. Repeat this step with the second well, transferring 10 µL to the third well, and so on, for as many serial dilutions as required. Plate the serial dilutions on an LB nutrient agar plate in 2 µL droplets, to ensure all droplets remain discrete. Incubate the LB Agar plates overnight at 37 °C and count visible CFUs. Calculate the number of CFUs per fly by counting the number of colonies present at the serial dilution where 0–40 CFUs are clearly visible. Then check the colony numbers in 10^-5^.

### Honeybee VPs and AMPs peptides generation

To generate the *UAS-*μ*elittin*, *UAS-Apamin*μ, *UAS-Apamin*^Δ*C*^, *UAS-tApamin*^Δ*C*^, *UAS-Apidaecin*, *UAS-Abaecin*, *UAS-Defendin-1*, *UAS-Defensin-2*, *UAS-Hymenoptaecin*, and *UAS-AMP5* driver in this study, peptide cDNAs were chemically synthesized with optimal *Drosophila* codon usage and with an optimal *Drosophila* Kozak translation initiation site upstream of the start methionine (CAAA)^73^. Encoded peptides are as follows: μ*elittin*, GIGAVLKVLTTGLPALISWIKRKRQQ; *Apamin*μ, CNCKAPETALCARRCQQH; *Apamin*^Δ*C*^, CNCKAPETALCA; *tApamin*^Δ*C*^, GAGFATPVTLALVPALLATFWSLLCNCKAPETALCA; *Apidaecin*, NNRPVYISQPRPPHPRL; *Abaecin*, YVPLPNVPQPGRRPFPT FPGQGPFNPKIKWPQGY; *Defendin-1*, VTCDLLSFKGQVNDSACAANCLSLGKAGGHCEKGVCICRKTSFKDLWDKRF; *Defensin-2*, SVPKVVYDGPIYELRQIEEENIEPDTELMDSNEPLLPLRHRRVTCDVLSWQSKWLSINHSACAIRCLAQRRKGGSCRNGVCICRK; *Hymenoptaecin*, QERGSIVIQGTKEGKSRPSLDIDYKQRVYDKNGMTGDAYGGLNIRPGQPSRQHAGFEFGKEYKNGFIKGQSEVQRGPGGRLSPYFGINGGFRF; *AMP5*, RDKKTLKIKMKLFI. These cDNAs were cloned into the pUAS-attB vector; For generation of transgenic *Drosophila*, Vectors was injected into the embryos of flies. The genetic construct was inserted into the attp40 site on chromosome II to generate transgenic flies using established techniques, a service conducted by Qidong Fungene Biotechnology Co., Ltd. (http://www.fungene.tech/).

### Immunostaining

After 5 days of eclosion, the *Drosophila* gut was taken from adult flies and fixed in 4% formaldehyde at room temperature for 30 minutes. The sample was than washed three times (5 minutes each) in 1% PBT and then blocked in 5% normal goat serum for 30 minutes. Subsequently, the sample was incubated overnight at 4℃ with primary antibodies in 1% PBT, followed by the addition of fluorophore-conjugated secondary antibodies for one hour at room temperature. For DAPI staining, the gut was incubated in DAPI containing 1%PBT for 10 minutes at room temperature, followed by three times wash. Finally, the gut was mounted on plates with an antifade mounting solution (Solarbio) for imaging purposes. Samples were imaged with Zeiss LSM880. Antibodies were used at the following dilutions: Chicken anti-GFP (1:500, Invitrogen), rat anti-deadpan (1:200, Abcam), Alexa-488 donkey anti-chicken (1:200, Jackson ImmunoResearch), Alexa-555 donkey anti-rat (1:200, Invitrogen), DAPI (1:1000, Invitrogen).

### Sample preparation and data analysis of 16S rRNA sequencing

The flies at 5-day age were collected under normal or infected conditions, each sample contains 25 flies with 3 replicas for each condition. Once collected in cryogenic tubes, they were frozen by liquid nitrogen and stored at -80 °C until measurement in the PCR by LC-BioTechnology Co., Ltd, HangZhou, Zhejiang Province, China. Samples were then sequenced on an Illumina NovaSeq platform according to the manufacturer’s recommendations, provided by LC-Bio.

### RNA extraction and cDNA synthesis

RNA was extracted from 50 preparations of 5-day-old females using the RNA isolation kit (Vazyme), following the manufacturer’s protocol. And first-strand cDNA was synthesized from 1μg of RNA template with random primers using SPARK script Ⅱ RT plus kit (SparkJade).

### Quantitative RT-PCR

The expression levels of *PGRP-SC2*, *DptA*, *AttC* in flies under normal or infected conditions were analyzed by quantitative real-time RT-PCR with SYBR Green qPCR MasterMix kit (Selleckchem). The primers of RT-PCR are *PGRP-SC2*, F:5’- GTTCTCGGCGTGACCATCAT-3’; R: 5’-TAGTTTCCAGCGGTGTGGTG-3’; *DptA*, F:5’-CACCGCAGTACCCACTCAAT-3’; R: 5’- AATCTCGTGGCGTCCATTGT-3’; *AttC*, F:5’-CGATGCCCGATTGGACCTAA-3’; R: 5’-ACTTGTTGTAGCCCAGGGTG-3’. qPCR reactions were performed in triplicate, and the specificity of each reaction was evaluated by dissociation curve analysis. Each experiment was replicated three times. PCR results were recorded as threshold cycle numbers (Ct). The fold change in the target gene expression, normalized to the expression of internal control gene (GAPDH) and relative to the expression at time point 0, was calculated using the 2 ^−ΔΔCT^ method as previously described^74^. The results are presented as the mean ± SD of three independent experiments.

### Locomotion assay

To detect and quantify the activity of flies, we have developed the Fly Trajectory Dynamics Tracking (FlyTrDT) software. This is an open-source, custom-written Python program that utilizes the free OpenCV machine vision library and the Python Qt library. The FlyTrDT software simultaneously records the trajectory information of each fly and calculates various indicators of the group at a certain period. For each frame acquired, the moving fly is segmented using the binarization function from the OpenCV library. Subsequently, a Gaussian blur and morphological closing and opening operations were performed on the extracted foreground pixels to consolidate detected features and reducing false positives and negatives. Finally, the extraction of fly outlines was achieved using the contour detection algorithm in the OpenCV library.

### Climbing assay

For climbing assay, we modified the conventional RING assay ^75^. In brief, 40-50 20- day-aged flies were placed in an empty vial and were tapped to the bottom of the tube. After tapping of flies, we recorded 10 seconds of video clip. This experiment was done five times with 5-minute intervals. With recorded video files, we captured the position of flies 10 seconds after tapping the vial. This captured image file was then loaded in ImageJ to perform particle analysis. For quantifying the location of flies inside a vial, we used the “analyze particles” function of ImageJ ^76^. The position of pixels was normalized by height of vial then only the particles above the midline (4 cm) of vial were counted.

### Lifespan Assay and Statistical Analysis

For lifespan analysis, we used conventional procedure as we described before ^77^. Briefly, 50 flies were aged by sex before being raised in typical 12 h light: 12 h dark cycles at either 25°C for each experimental objective. The number of dead flies was recorded every two to three days. Every three to four days, the surviving flies were transferred to fresh vials.

### Single-fly sleep and circadian rhythm recording

96-well white Microfluor 2 plates (Fishier) with 400 μl of food (5% sucrose and 1% agar) were loaded with adult male flies (aged 3–5 days). Flies were entrained to the 12 h:12 h LD cycles for four days at 25 °C to record sleep behavior, then changed to constant darkness for 5-6 days to record circadian rhythms in the absence of light inputs. The fly movement was monitored using a camera at 10s intervals, and the data were then used by the sleep and circadian analysis program SCAMP to analyze sleep and circadian rhythm ^78–80^. It calculates activity by shifting the position of *Drosophila* every 10 seconds and calculates sleep using the standard definition (*Drosophila* is recorded as asleep if it remains motionless for at least 5 minutes).

### Smurf assay

For Smurf assay, 3% (v/v) Food Blue No.1 aluminum lake (Aladdin, F336821) was mixed with normal food, and flies under different conditions were imaged on Olympus SZ61 microscope after a 6-hour feeding or 20-hour feeding^81^. For flies under sleep deprivation, a vortex machine (CHANGZHOU ENPEI INSTRUMENT MANUFACTURING CO., LTD., NY-5SX) was applied with a routine of 2-second 1500 rpm vortex following a one-minute rest^82^, flies were transferred in tubes containing normal food and went through a 24-hour sleep deprivation. For DSS feeding, 3% DSS (Coolaber, 9011-18-1) was mixed with blue dye food and flies were transferred and fed overnight^83^.

### Phenol red pH test

Flies were transferred to food containing 0.2% Phenol Red (Macklin, P6066)^84^, and fed for at least 6 hours. Gut was dissected in PBS and imaged under microscope.

### DHE staining for ROS detection

The procedure was modified based on protocol published previously, and was briefly described as follows^85^. Fly gut was dissected and incubated in Schneider ’s Fruit Fly culture medium (with Glutamine) (VivaCell). Make a 30mM stock solution of DHE (Invitrogen, D11347) in anhydrous DMSO (Sigma-Aldrich, cat. no. 276855), dissolve 1 µl of the dye in 1ml of schneiders medium to give a final concentration of approximately 30uM. Vortex to evenly disperse the dye. Incubate the gut with the dye for 7 minutes in a dark chamber, followed with three 5-minute washes in schneiders medium. And the gut was mounted on plates with an antifade mounting solution for imaging purposes after DAPI staining as described before.

### Honeybee infection assay

The European honeybees, *Apis mellifera*, in this study were kindly provided and maintained by Dr. Fangyong Ning (Northeast Agricultural University, Harbin, China). For oral administration, the procedure was modified based on previous study, and was described as follow^86^. Bees were anesthetized on ice and placed in 1.5 ml tubes with the assistance of paper tapes for fixation. Bees were recovered from anesthesia at room temperature and fed 10 µl of each sample. For control groups, bees were administrated with 1 mol/L sucrose and for bacterial infection group, bees were fed with bacteria pellet resuspended by 1 mol/L sucrose. As for apamin administration, apamin powder (Chemstan, 24345-16-2) was dissolved into 1 mol/L sucrose at a high concentration of 0.01mg/mL, and 100 µL of high concentration apamin was transferred to 1.5 ml tube, adding 900 µL 1 mol/L sucrose creating a medium concentration. Repeat this step to create low concentration apamin, and apamin at various concentrations were applied to honeybees using a micropipette. Treated bees were maintained in a plastic box and fed with small honey comb at 25°C. Honeybees were judged to be dead and recorded if no motion was detected in any body parts.

### Statistics

All analysis was done in GraphPad (Prism9). Besides traditional *t*-test for statistical analysis, we added estimation statistics for all two group comparing graphs in bacterial infection assay, immunostaining comparison and Smurf assay data. To compare the survival curves of each genotype, the data was analyzed by log-rank (Mantel-Cox) test. In climbing assays, mean values were compared by one-way ANOVA, each figure shows the mean ± standard deviation (SD) (\**\*\*\* = p<0.0001, *** = p < 0.001, ** = p < 0.01, * = p < 0.05*, n.s. stands for non-significant differences).

## Data and code availability

- All data reported in this paper will be shared by the lead contact upon request.
- This paper does not report original code. The URL of the codes used in this paper are listed in the key resources table.
- Any additional information required to reanalyze the data in this paper is available from the lead contact upon request.

## SUPPLEMENTAL FIGURE TITLES AND LEGENDS

**Fig.S1: The chemical structure of apamin, the antimicrobial effect, survival curve and locomotion of flies expressing different honeybee VPs.**

**a,** Chemical structure of apamin and mature apamin amino acids sequence. **b,** Infectious dose of male flies expressing mature format melittin by *tub-GAL4* following a 12-hour orally feeding of *P. aeruginosa* culture. CFU stands for Colony- Forming Unit. DBMs represent difference between means, which is a statistical measure that quantifies the average discrepancy between two groups. (n=10) **c,** The climbing ability of female flies expressing mature format melittin by *tub-GAL4*. (n=5) **d,** The climbing ability of male flies expressing mature format melittin by *tub-GAL4*. (n=5) **e,** Infectious dose of male flies expressing Apamin^ΔC^ via *tub-GAL4*. (n=10) **f,** Infectious dose of female flies expressing Apamin^ΔC^ by *tub-GAL4*, da-GAL4 compared with control. (n=10) **g,** The trajectory percentage of female Apamin^ΔC^ and tApamin^ΔC^ flies compared with control flies. (n=49) **h,** The survival curve of control male flies and male flies expressing Apamin^ΔC^ and tApamin^ΔC^ via by *tub-GAL4* driver. (n=50)

**Fig.S2: 16S rRNA sequencing results of control flies and flies expressing tApamin^ΔC^.**

**a,** Heatmap of the top 31 significant genus (P value < 0.05, in the comparison of infected and uninfected, tApamin^ΔC^ versus control; n = 3 for each condition). **b,** Dendrogram and histogram of the bacterial composition at the genus level. **c,** Bacterial KEGG pathway prediction by PICRUSt2. Black bar: Foldchange of relative abundance; white bar: −log10P values for each KO term.

**Fig.S3: Tissue screening of antimicrobial effects in tApaminΔC expressing male flies and gut environment changes displayed.**

**a-h,** Infectious dose between male flies expressing tApamin^ΔC^ via **a,** *tub-GAL4* and *Hml-Gal4*, **b,** *Pxn-GAL4*, **c,** *Cg-GAL4*, **d,** *3.1Lsp2-GAL4*, **e,** *Drip-GAL4*, **f,** *Akh- GAL4*, **g,** *Pros-GAL4*, **h,** *esg-GAL4*, following a 12-hour orally feeding of *P. aeruginosa* culture. (n=10) **i,** Infectious dose between female flies expressing tApamin^ΔC^ via *tub-GAL4* and *Mhc-Gal4*. (n=10) **j,** Infectious dose between male flies expressing tApamin^ΔC^ via *tub-GAL4* and *Mef2-Gal4*. (n=10) **k, l,** The ISCs in full female fly gut between normal condition **(k)** and tApamin^ΔC^ expression flies **(l)** driven by *esg-GAL4* with *UAS-GFP*, fly guts were immunostained with anti-GFP (green), and DAPI (blue). **m-p,** The Dpn-positive neuroblast cells and mitochondria (*UAS- mitoGFP*) in full gut of female flies in normal **(m)** and tApamin^ΔC^ expressing conditions **(n)** by *tub-GAL4*, and the rectal pupillae part respectively **(o,p)**. Fly guts were immunostained with anti-Deadpan (red), anti-GFP (green), and DAPI (blue). **q,r,** DHE staining of ROS level in full female fly gut between control **(q)** and tApamin^ΔC^ expressing flies **(r)** driven by *tub-GAL4*. Fly guts were stained with DHE (red) and DAPI (blue) (repeated for 3 times).

**Fig.S4: Locomotion of neuronal Apamin^ΔC^ expressing fly, sleep pattern of pan- expressed Apamin^ΔC^ fly and Smurf related results induced by stress.**

**a,** The trajectory length, **b,** trajectory area, **c,** rotation times, **d,** stop counts, and **e,** trajectory diagram of female flies expressing Apamin^ΔC^ by *nSyb-GAL4* (n=48) compared with controls (n=50). **f-o,** Sleep profiles (average proportion of time spent sleeping in consecutive 30-min segments during a 24-h LD cycle) and quantification of female flies expressing Apamin^ΔC^ expression by *tub-GAL4* driver (n=11 for controls, n=12 for Apamin^ΔC^ expressing flies) **(f)**. Quantification of sleep durations for Apamin^ΔC^ expression flies **(g-i)**. Quantification of sleep episodes for Apamin^ΔC^ expression flies **(j-l)**. Quantification of sleep latency for Apamin^ΔC^ expressing flies **(m-o)**. **p,** The mean intensity of Smurf assay flies expressing tApamin^ΔC^ compared with controls after 24 hours sleep deprivation and 6 hours dye feeding. The experiment was repeated three times and one representative figure was shown of each condition. **q,** The counts of facets on plugs containing flies treated with a 3% DSS feeding with blue dye. The experiment was repeated three times and one representative figure was shown of each condition. **r,** The counts of facets on tube wall of flies treated with a 3% DSS feeding with blue dye. The experiment was repeated three times and one representative figure was shown of each condition (repeated for 3 times).

**Fig.S5: Possible immune genes and AMPs related, and survival of honeybees applied to apamin in different concentrations.**

**a,** Infectious dose of male flies expressing Apamin^ΔC^ and tApamin^ΔC^ by *tub-GAL4*, with a knockdown of *Relish* compared with control. (n=10) **b,** Infectious dose (n=10) and **c,** Climbing ability (n=5) of controls and male flies expressing Apidaecin by *tub- GAL4* following a 12-hour orally feeding of *P. aeruginosa* culture. **d,** Infectious dose of controls and male flies expressing Abaecin by *tub-GAL4*. (n=10) **e,** Climbing ability of controls and female flies expressing Abaecin via *tub-GAL4*. (n=5) **f,** Climbing ability of controls and male flies expressing Abaecin via *tub-GAL4*. (n=5) **g-j,** Infectious dose of controls and male flies expressing Defendin-1 **(g)**, Defensin-2 **(h)**, Hymenoptaecin **(i)**, and AMP5 **(j)** by *tub-GAL4* following a 12-hour orally feeding of *P. aeruginosa* culture. (n=10) k, The survival curve of honeybees infected orally by *P. aeruginosa*, following sucrose administration as control and three different concentrations of apamin administration. Control (black) (n=9) compared with low concentration apamin (yellow) (n=9), medium concentration apamin (orange) (n=10), and high concentration apamin (red) (n=10) administration.

**Movie.S1: Honeybees after apamin administration.**

**Movie.S2: Honeybee apamin and bacteria administration.**

