## Supplementary figures and images for "Investigating the antimicrobial and immunomodulatory effects of honeybee venom peptide apamin in the *Drosophila* genetic platform"

### Supplemental Figure 1

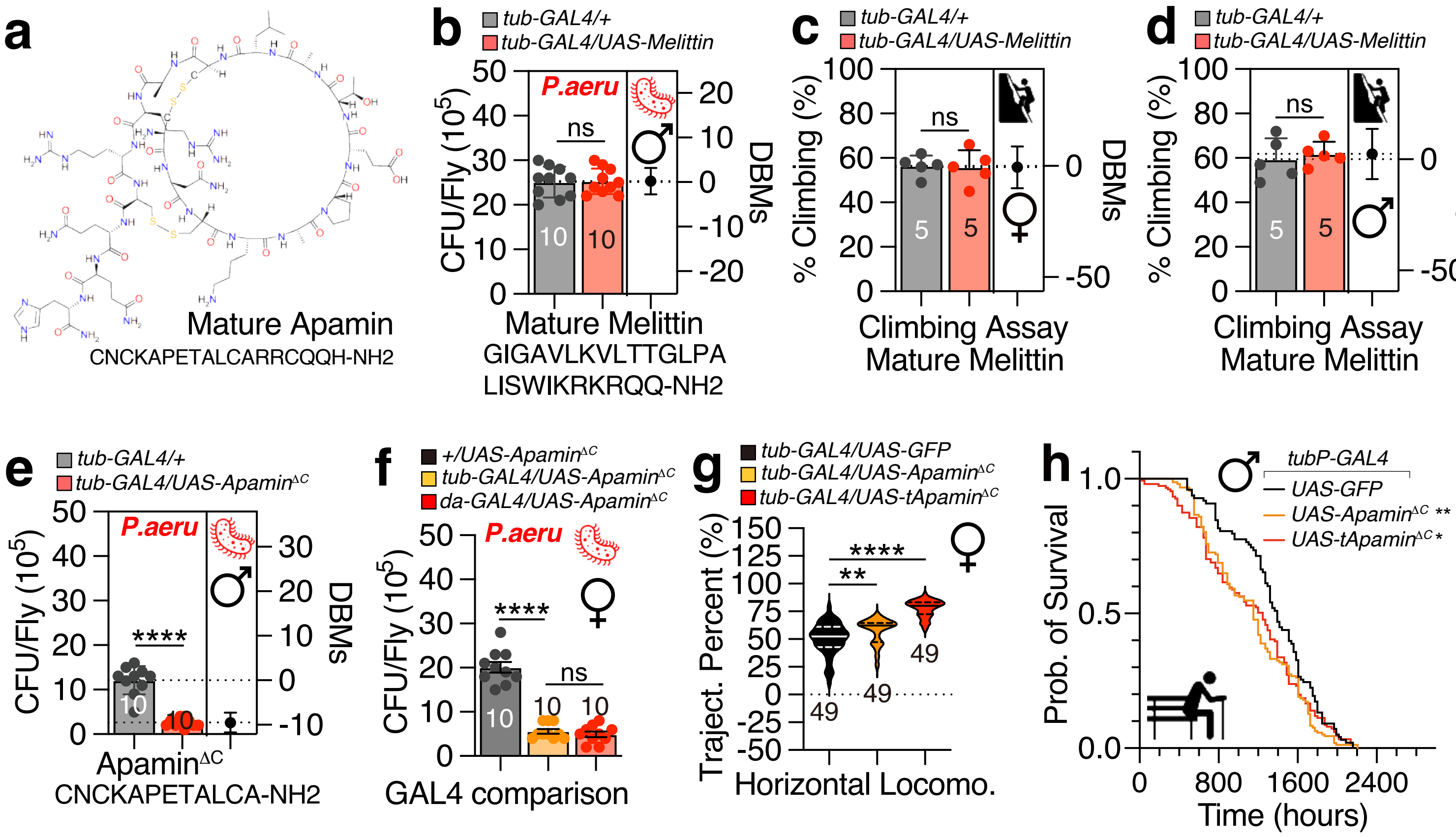


**Apamin, Fig.S1**

### Supplemental Figure 2

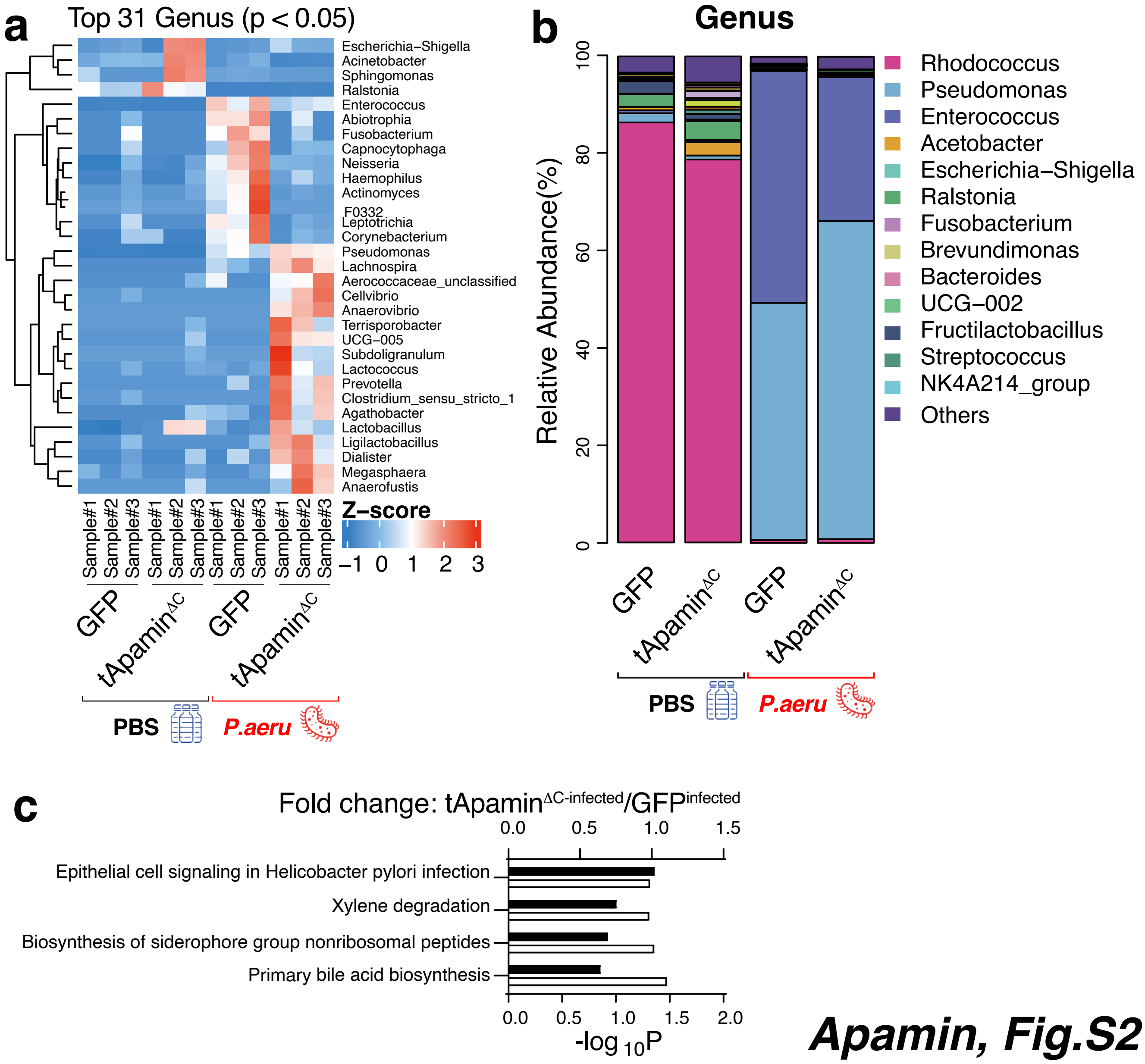


***Apamin, Fig.S2***

### Supplemental Figure 3

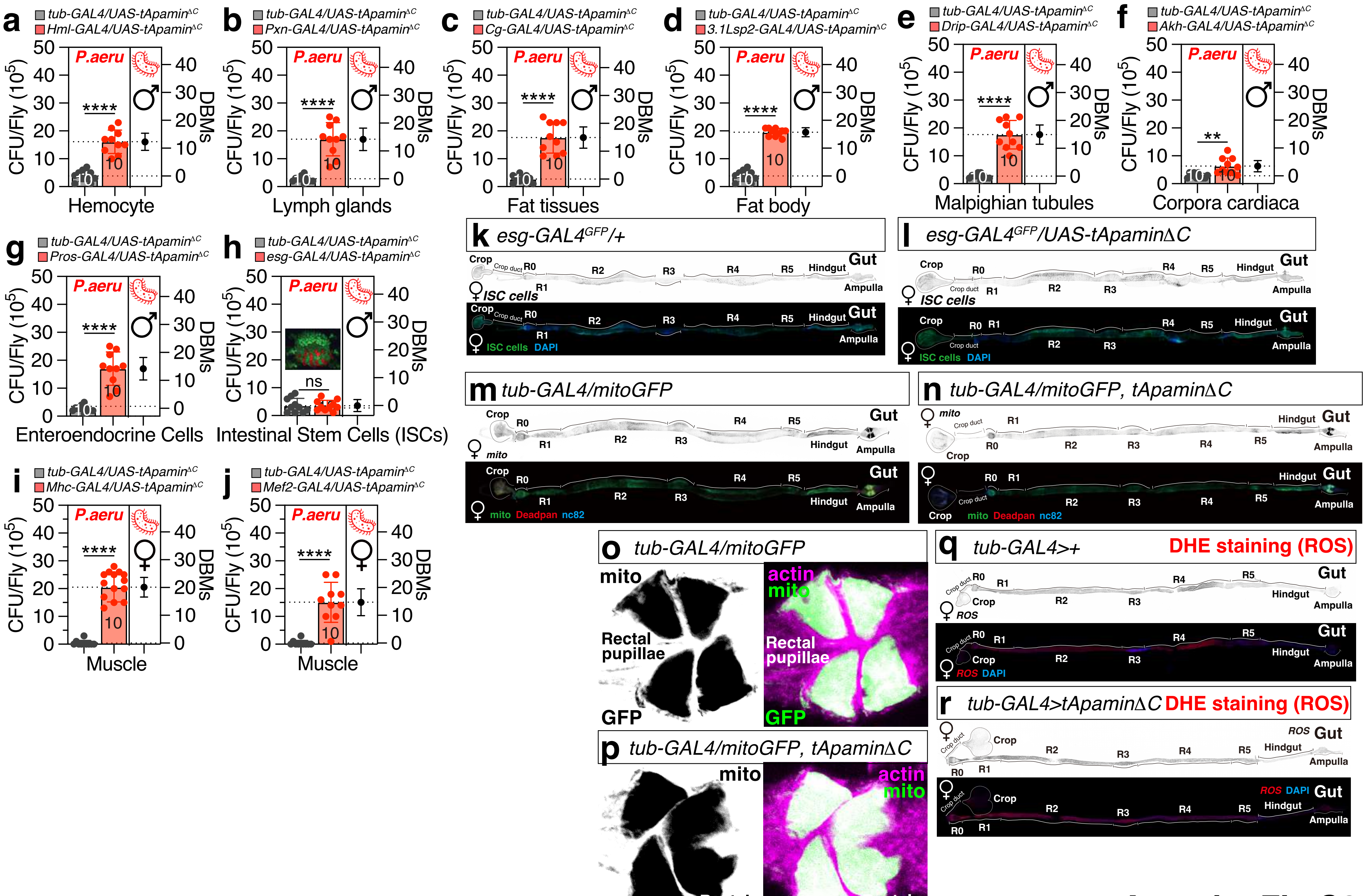


**Apamin, Fig.S3**

### Supplemental Figure 4

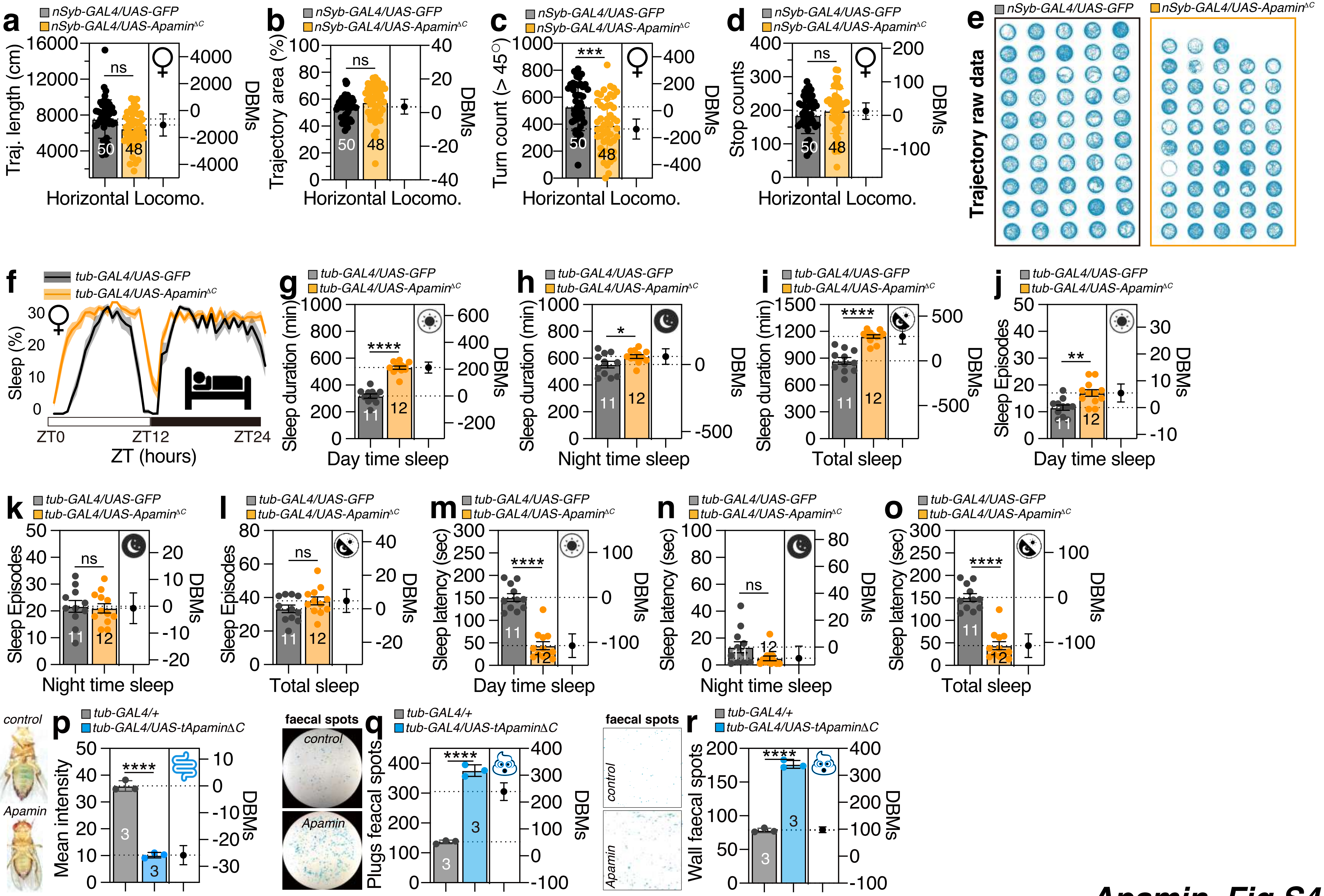


**Apamin, Fig.S4**

### Supplemental Figure 5

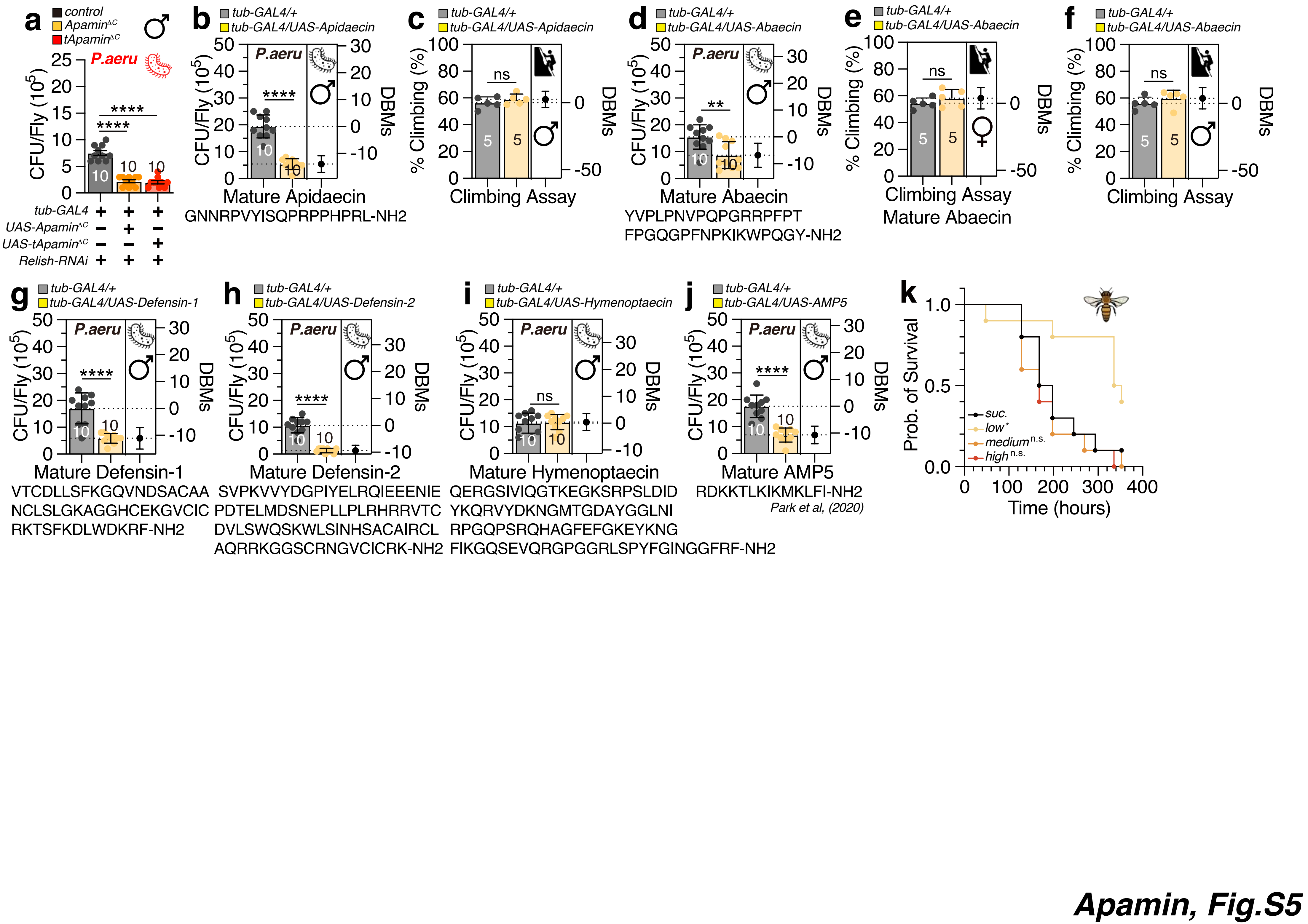
